# Semi-automated curation and manual addition of sequences to build reliable and extensive reference databases for ITS2 vascular plant DNA (meta-)barcoding

**DOI:** 10.1101/2023.06.12.544582

**Authors:** Andreia Quaresma, Carlos Ariel Yadró Garcia, José Rufino, Mónica Honrado, Joana Amaral, Robert Brodschneider, Valters Brusbardis, Kristina Gratzer, Fani Hatjina, Ole Kilpinen, Marco Pietropaoli, Ivo Roessink, Jozef van der Steen, Flemming Vejsnæs, M. Alice Pinto, Alexander Keller

**Affiliations:** Centro de Investigação de Montanha (CIMO), Instituto Politécnico de Bragança, Campus de Santa Apolónia, 5300-253 Bragança, Portugal; Laboratório Associado para a Sustentabilidade e Tecnologia em Regiões de Montanha (SusTEC), Instituto Politécnico de Bragança, Campus de Santa Apolónia, 5300-253 Bragança, Portugal; Departamento de Biologia, Faculdade de Ciências da Universidade do Porto, Rua do Campo Alegre, S/N, Edifício FC4, 4169-007, Porto, Portugal; CIBIO, Centro de Investigação em Biodiversidade e Recursos Genéticos, InBIO Laboratório Associado, Campus de Vairão, Universidade do Porto, 4485-661 Vairão, Vila do Conde, Portugal; BIOPOLIS Program in Genomics, Biodiversity and Land Planning, CIBIO, Campus de Vairão, 4485-661 Vairão, Vila do Conde, Portugal; Institute of Biology, University of Graz, Universitätsplatz 2, 8010 Graz, Austria; Latvian Beekeepers’ Association (LBA), Rigas iela 22, LV-3004 Jelgava, Latvia; Ellinikos Georgikos Organismos DIMITRA (ELGO- DIMITRA), Kourtidou 56-58, GR-11145 Athina, Greece; Danish Beekeepers Association (DBF), Fulbyvej 15, DK-4180 Sorø, Denmark; Istituto Zooprofilattico Sperimentale del Lazio e della Toscana “M.Aleandri” (IZSLT), Via Appia Nuova 1411, IT-00178 Roma, Italy; Wageningen Environmental Research (WENR), Droevendaalsesteeg 3, NL-6708 PB Wageningen, The Netherlands; Research Centre in Digitalization and Intelligent Robotics (CeDRI), Instituto Politécnico de Bragança, Bragança, Portugal; Alveus AB Consultancy, Kerkstraat 96, NL-5061 EL Oisterwijk, The Netherlands; Organismic and Cellular Interactions, Biocenter, Faculty of Biology, Ludwig-Maximilians-Universität München, Martinsried, Germany

## Abstract

With the breakthrough of DNA (meta)-barcoding, it soon became clear that one of the most critical steps for accurate taxonomic identification is to have an accurate DNA reference database for the DNA barcode marker of choice. Therefore, developing such a database has been a long-term ambition, especially in the Viridiplantae kingdom. Typically, reference databases are constructed from marker sequences downloaded from general public databases, which can carry taxonomic and other relevant errors. Herein, we constructed a curated (i) global database, (ii) European crop database, and (iii) 27 databases for the EU countries for the ITS2 barcoding marker of vascular plants. To that end, we first developed a pipeline script that entails (i) an automated curation stage comprising five filters, (ii) manual taxonomic correction for misclassified taxa, and (iii) manual addition of newly sequenced species. The pipeline allows easy updating of the curated global database. With this approach, 13% of the sequences, corresponding to 7% of species, originally imported from GenBank were discarded. Further, 259 sequences were newly added to the curated global database, which now comprises 307,977 sequences of 111,382 plant species.

## Background & Summary

DNA barcoding, a concept put forward by Hebert et al.^1^ in 2003, was developed to facilitate species identification using molecular methods. DNA barcoding standardizes the taxonomic identification of organisms based on well-established short genomic regions that have high interspecific and low intraspecific variability. By definition, a DNA barcoding marker must be universal, reliable, and show good discriminatory power at the species level^2^. For animals and fungi, the mitochondrial cytochrome c oxidase I gene (COI)^1^ and the internal transcribed spacer (ITS) region^3^, respectively, have been defined and accepted by the scientific community as the genomic regions that fulfil these criteria. However, in the *Viridiplantae* kingdom, there is no single barcoding marker that satisfies all of those criteria, and several markers in the mitochondrial, chloroplastidial, and nuclear genomes have been under dispute^4-6^. Finally, four DNA barcoding markers have been agreed upon for taxonomic identification of plants, including the chloroplastidial regions rbcL, matK, and trnH-psbA, as well as the nuclear internal transcribed spacer (ITS) region of the ribosome, particularly the ITS2 region^2,7,8^.

The emergence of high-throughput sequencing (HTS) techniques is tightly linked to the recent burst of DNA metabarcoding studies^9,10^, which have used one or more of the four markers for taxonomic identification. DNA metabarcoding is a powerful approach for resolving mixed-species samples or environmental DNA (eDNA)^11^ at large spatial scales, with multiple applications in the fields of ecology, taxonomy, evolution, and conservation^11,12^ for a wide array of organisms. In plants, DNA metabarcoding has been applied in the authentication of herbal teas^13-15^, determining herbivore diets^16-19^, unravelling plant-pollinator interactions^20-23^, identifying botanical origin of honey^24-26^, monitoring allergy-related airborne pollen sources^27,28^, assessing biodiversity^29-32^, or even in forensic analysis^33^. These studies have either employed single DNA marker or their combinations, with most relying on rbcL and/or ITS2^34-36^ (Fig 1). ITS2 has been increasingly popular (Fig 2), due to its better taxonomic discriminatory capabilities^2^ and the higher number of sequences available in GenBank^37^ (Table 1) as compared with the other three plant barcoding markers.

**Fig. 1.**
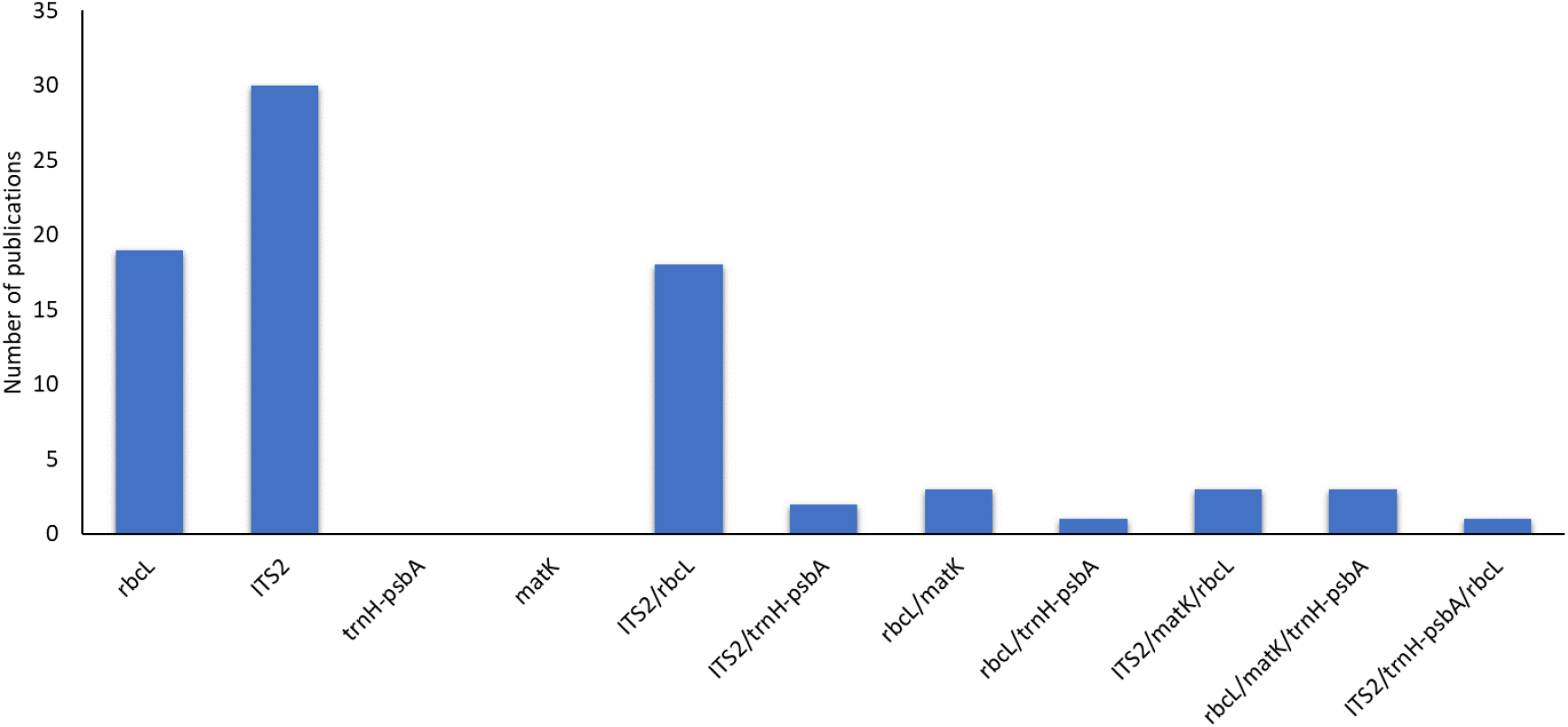
Number of publications for each of the *Viridiplantae* DNA barcoding markers, used singly or in combination in DNA metabarcoding studies. Information retrieved from Web of Science, accessed on 22 February 2023, using the following search strings: “ITS2 (All Fields) AND metabarcoding (All Fields) NOT fung* (All Fields) NOT diatoms (All Fields)”; “rbcL (All Fields) AND metabarcoding (All Fields) NOT fung* (All Fields) NOT diatoms (All Fields)”; “trnH-psbA OR psbA-trnH (All Fields) AND metabarcoding (All Fields) NOT fung* (All Fields) NOT diatoms (All Fields)”; “matK (All Fields) AND metabarcoding (All Fields) NOT fung* (All Fields) NOT diatoms (All Fields)”. The retrievals were manually verified and the publications that were not targeted to plants were ignored.

**Fig. 2.**
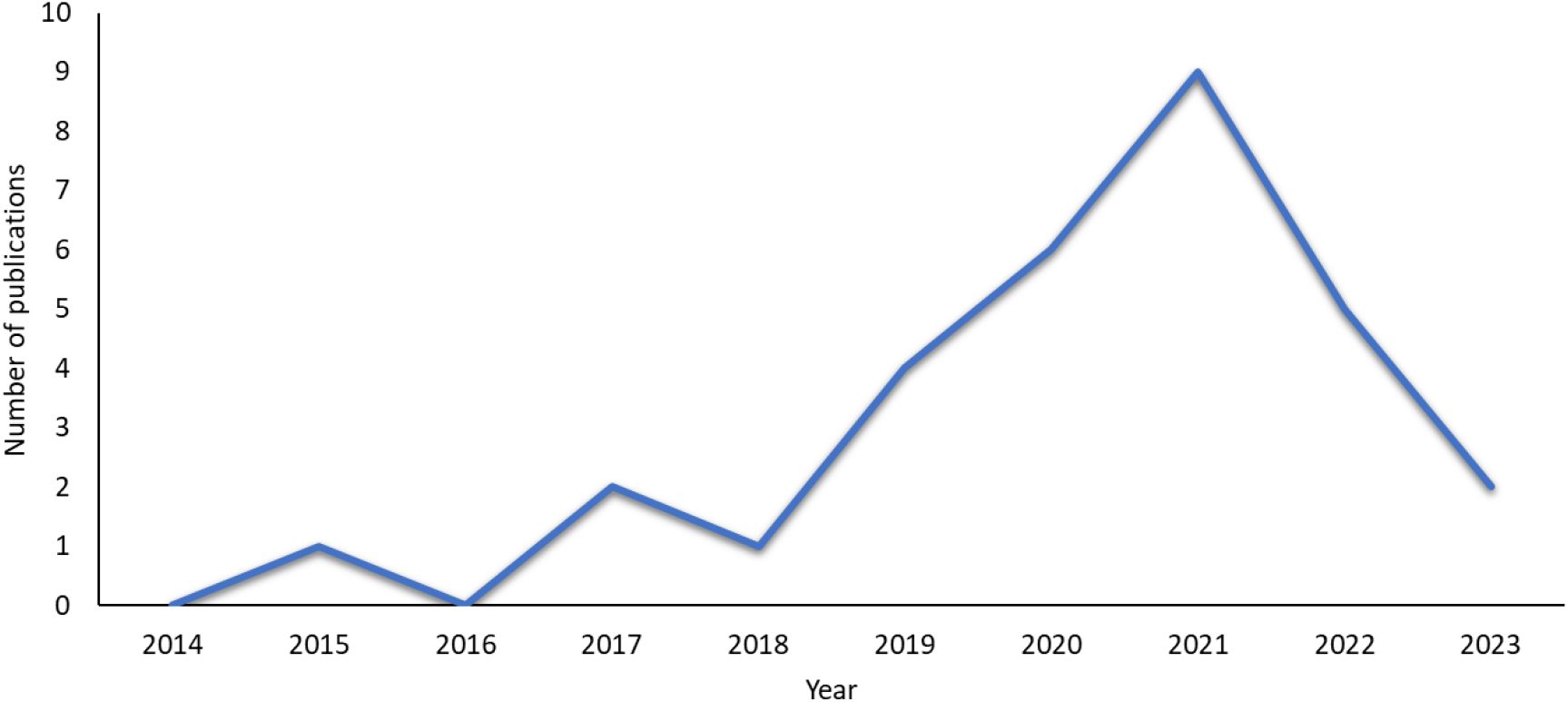
Number of publications on ITS2 metabarcoding in the last ten years. Information retrieved from Web of Science, accessed on 22 February 2023, using the search string “ITS2 (All Fields) AND metabarcoding (All Fields) NOT fung* (All Fields) NOT diatoms (All Fields)”.

**Table 1.**
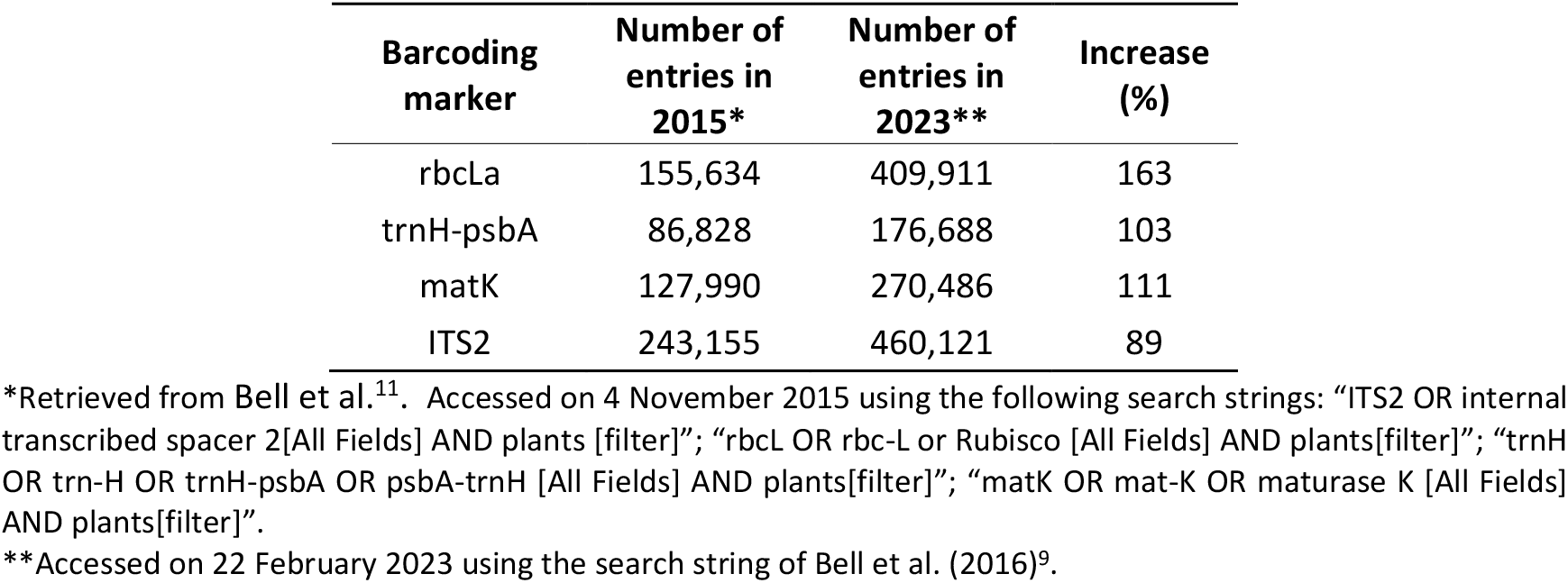
Number of sequences available in GenBank for each of the *Viridiplantae* DNA barcoding marker in 2015 and 2023, and corresponding increase rate during this period.

Botanical identification of mixed-species samples by DNA metabarcoding entails their laboratorial processing to obtain the sequence reads with the HTS technology. The millions of reads generated by the HTS run are then classified against sequences of known taxonomic origin, which are typically compiled into a reference database constructed using own sequences or such retrieved from GenBank or other public databases. The quality of identification depends on the quality and completeness of the reference database built for the target barcoding marker, which in turn is determined by the breadth and size of the database (number of taxa and number of sequences per taxon) as well as by the taxonomic accuracy of the compiled sequences^20,38,39^. Many plant studies have relied on sequence data directly retrieved from GenBank for identifying unknown samples^24,40-43^. The problem with this approach is that sequences deposited in GenBank are not rigorously checked for taxonomic mistakes and other inconsistencies that might affect barcoding purposes. Erroneous records are common and can for example be due to fungi, deposited on the surface or inside of plant tissues and sequenced instead of the targeted plant, or to plants that were morphologically misidentified^38^. This results in inaccurate classifications using direct hit methods (e.g., VSEARCH^44^, USEARCH^45^, BLAST^46^), and also in poor models for hierarchical classifications (e.g., RDPclassifier^47^, SINTAX^48^).

Construction of high-quality reference databases for plants has been sought over the years, and several attempts have been made, specifically for ITS2 and rbcL. The first ITS2 reference database was released in 2006^49^ for different kingdoms. This database underwent several updates until 2015^50^. In the same year, Sickel et al.^51^ built the first *Viridiplantae* ITS2 database from the earlier multi-kingdom one, which has been used in several plant metabarcoding studies^34,20,52,53^. However, due to the ever-increasing number of sequences deposited in GenBank, this database soon became outdated (Table 1). In 2017, Bell et al.^52^ developed a rbcL database, which was combined with the existing ITS2 *Viridiplantae*^*51*^, for species level identification in angiosperms. This rbcL database was last updated in 2021, at the same time that a new ITS2 database for *Magnoliopsida* was developed by the same group^54^. In 2019, Curd et al.^55^ developed the ANACAPA toolkit, which comprises a module to generate custom reference databases for any marker. In 2020, Banchi et al.^38^ published an ITS database, named PLANiTS, that includes datasets for ITS, ITS1 and ITS2. In addition, these authors developed a script that performs a species identity check on the sequences downloaded from GenBank, although it is a QIIME2 based script. Also in 2020, Richardson et al.^56^, developed the toolkit MetaCurator, which generates reference databases dedicated to taxonomically informative genetic markers, while Keller et al.^39^ developed BCdatabaser, a tool that allows generating generic databases of any marker by linking sequences and taxonomic information retrieved from GenBank. In 2022, Dubois et al.^12^ developed a workflow that allows the building of plant reference databases dedicated to ITS2 and rbcL. However, this workflow can only be used on the QIIME2 platform.

The first developed databases were static and, therefore, easily outdated due to the rapid flow of new sequences being deposited in GenBank (Table 1). Moreover, most of the available databases are global-scale, which may lead to taxa misidentifications because of sequence conflicts originating from misidentified GenBank sequences or even from polyphyletic species. Accordingly, it might be helpful to have a database tailored for the geographical area under analysis, as a way of reducing the identification error by including only the extant flora, therefore minimizing the detection of unlikely taxa^57^. Complementary to this, it is also important to have user-friendly tools that automatically perform the generation and curation of a reliable and updatable reference databases. Currently, most of the available tools require some level of user bioinformatics expertise or lack a good curation method for handling the problem of misidentified GenBank sequences. For instance, BCdatabaser^39^ is a friendly tool as it entails a single command to produce a taxonomy-linked *fasta* file, which can be used by several taxonomic classifiers. However, it lacks a curation method, and it includes the download of non-target sequences incorrectly annotated in GenBank^12^ (e.g., rbcl sequences that are identified as ITS2).

In this context, the goal of this study was to provide curated databases for ITS2 (meta)-barcoding, and a reproducible, public, and pipeline-based workflow that is compatible with other custom databases. The script was designed to be applied after using BCdatabaser^39^ or similar workflows that generate taxonomically linked *fasta* files. The workflow consists of three main stages: (i) automated curation of the downloaded sequences that accounts for five major problems detected in sequences deposited in GenBank (fungal sequences identified as vascular plants, *Chlorophyta* sequences, non-target sequences, incomplete taxonomies, and erroneous taxonomy annotation); (ii) a manual taxonomic correction option for misidentified taxa; and (iii) the addition of custom sequenced species to conform with the common syntax of the database. Using this workflow, we generated an ITS2 reference database that comprises worldwide vascular plant taxa, as well as individual subsets of this database for each of the 27 countries of the European Union, and a reference database for European crops.

## Methods

### Curation pipeline-based workflow

The pipeline-based workflow comprises three independent stages for generating a more accurate reference databases: (i) automated curation, (ii) manual list curation, and (iii) manual sequence addition (Fig. 3). These can be performed singly or in conjunction, depending on the user’s needs. The pipeline script is publicly available at GitHub (https://github.com/chiras/database-curation) and has as dependencies the also publicly available software tools R^58^, SeqFilter v2.1.10^59^ (https://github.com/BioInf-Wuerzburg/SeqFilter), and VSEARCH v2.18.0^44^ (https://github.com/torognes/vsearch). It is designed to start after the point of pulling reference sequences from GenBank with BCdatabaser (or equivalent tools), or from other public sources that follow the same syntax needed for a variety of classifiers (https://molbiodiv.github.io/bcdatabaser/output.html). The pipeline was executed successfully on the bash command line of Ubuntu 20.04.6 and Mac OSX 12.3.

**Fig. 3.**
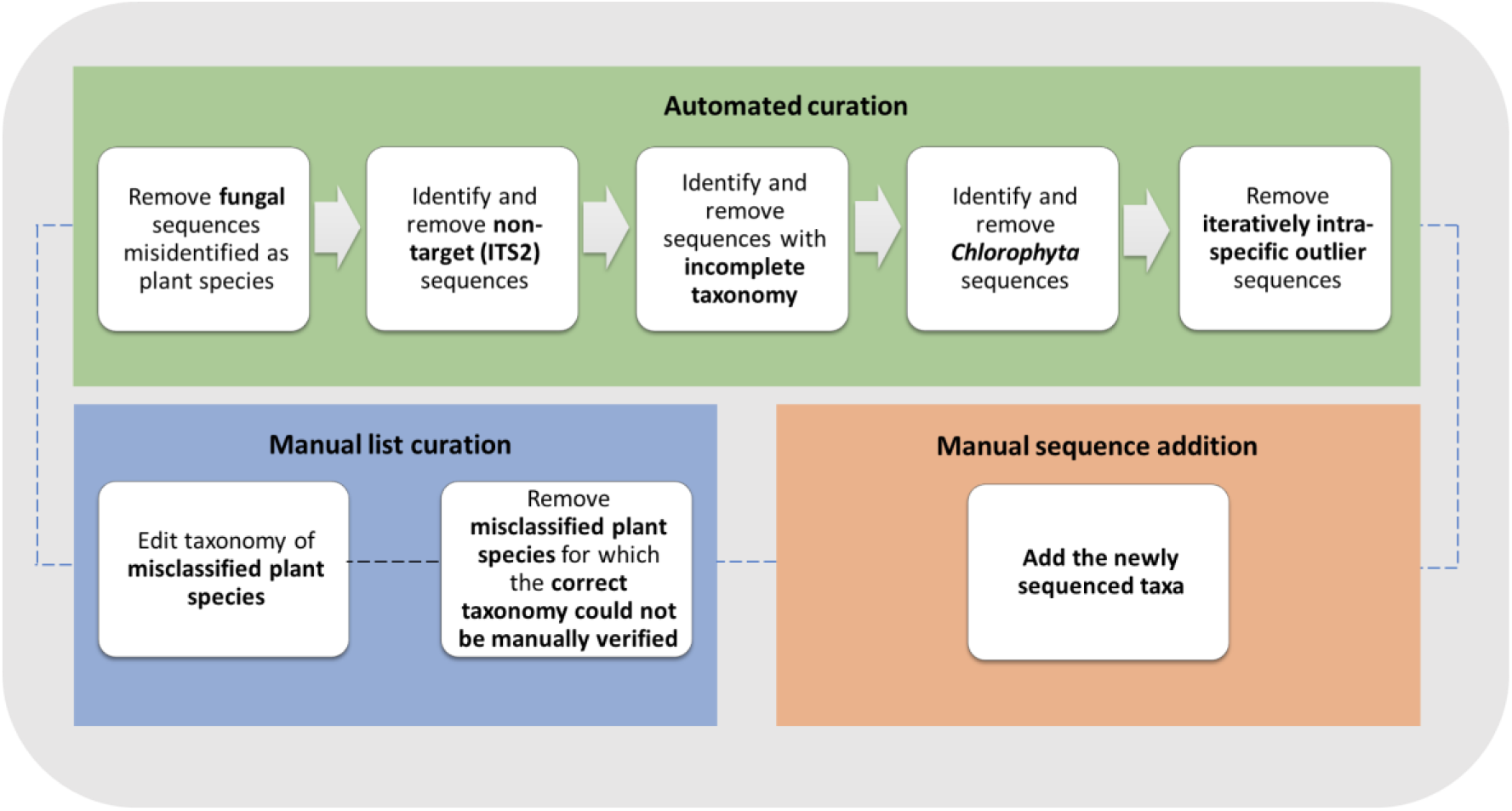
Schematic representation of the curation pipeline. The components ‘Automated curation’, ‘Manual list curation’, and ‘Manual sequence addition’ can be used singly or in conjunction.

#### Automated curation

The automated curation is the most important stages of the pipeline-based workflow. Five major cleaning steps are implemented during curation (Fig. 3):

i. The first filter identifies fungal sequences and removes them. These are identified by using a hierarchical classification with the *sintax*^*48*^ command from VSEARCH against the RDP curated fungal ITS database^60^, with a cut-off of 0.90;
ii. The second filter performs the removal of non-ITS2 (non-target) sequences. For this, we manually created a preliminary ITS2 reference database of selected trustworthy sequences representing all vascular plant families from the ITS2 database^50^. In the automated curation, the command *usearch_global* by VSEARCH is used to identify only vascular plant sequences with an identity threshold of 70%;
iii. The third filter checks for incomplete taxonomy entries in the metadata and removes such entries as they are not suitable for barcoding purposes and might interfere with finding better resolved references;
iv. The fourth filter removes all the sequences that are classified as *Chlorophyta* as our intention was to create a reliable vascular plant database. Wrong annotations of *Chlorophyta* sequences can also interfere with vascular plant identification;
v. The fifth filter applies a deterministic assessment of intraspecific variability for the respective database on-the-fly. However, this filter is only applied to species that are represented by more than four sequences. The database is hereby split into subsets for each plant species, and, for each separate species database, pairwise all-against-all global alignments are performed with *allpairs_global* from VSEARCH. An iterative R script increases a drop-out threshold for each species in steps of 50%, 75%, 80%, 85%, 90%, 92.5%, 95%, and 97%, removing sequences that have a lower median identity to all other sequences of the species than the threshold, but only while a threshold is given that removes less than 50% of the remaining sequences per species. This 50% threshold is a balanced trade-off between removing taxa with wrong GenBank taxonomic assignments and retaining sequences that are still within expected intraspecific variability (see ‘Assessment of intraspecific variability’ section for further details).

#### Manual list curation

The manual list curation is intended to serve as a community-driven approach. Scientists that spot erroneous GenBank entries that are not identified by the automated curation are invited to add a simple tabular text file to our GitHub repository. Based on these text files, researchers curating a database can choose to use or discard manual curations from different contributors. The text file format is kept as simple as possible, and examples are given in the code repository:

~~~
NCBIAccessionNumber;Wrong_ScientificName;Corrected_ScientificName;Curator_N ame.
~~~

If the file specifies the `Corrected_ScientificName`, the script will proceed to correct the taxonomy in the reference database. The field can be left empty as well, indicating that the curator is sure that this is a wrong taxonomic metadata and yet unsure about the correct identification, which will result in the sequences being removed from the database.

#### Manual sequence addition

The manual addition allows users to add own generated sequences to the reference database, and automating the gathering of taxonomic metadata and formatting. This is a tedious step, especially when many sequences are added. The requirement for this step is the provision of common *fasta* files with the species name as the header. Examples are provided in the GitHub repository.

#### Global database subdivision

The subdivision of the global database allows the user to reduce the number of species from the global reference database to a local reference database that contains a geographically delimited number of species. For this step, it is required to provide a list of the intended local flora in a csv file format.

### Application of the pipeline, curation of ITS2 databases, and enrichment with new sequences

#### Data acquisition

A *Viridiplantae* ITS2 reference database, hereafter called “global database” was created on 17 January of 2023 using BCdatabaser^39^. This dataset comprises a maximum of 25 sequences per species of the *Viridiplantae* kingdom, with a length between 100-2,000 bp. Across the study, we found that crop species represent a special case for barcoding purposes because they show a high intraspecific variability of the ITS2 region, often due to hybridizations or other genomic intervention (e.g.: *Brassica* and *Malus*). Therefore, we considered that there was an additional need for developing a reference database only for the European crops, which is further referred to as the “crop database”. This dataset was generated in the same way, but instead of 25 sequences, a maximum of 100 ITS2 sequences per species was downloaded from GenBank to account for a higher representation of intraspecific variability.

In addition, 536 leaf samples representing 322 species, selected from expert knowledge as important pollen sources for the honey bee (*Apis mellifera*), collected from nine European countries (Austria, Denmark, France, Greece, Italy, Latvia, The Netherlands, Norway, Portugal) were further sequenced for the ITS2 region, aiming for manual addition into the database (Table 2). These species were missing or underrepresented in the initial global database. The leaves were cut in small pieces and transferred to a 2.0 ml screwcap tube with two 3 mm zirconia beads. After being grounded in a Precellys 24 tissue homogeniser (Bertin Instruments), the DNA was extracted with the Macherey-Nagel NucleoSpin Plant II Kit, according to the manufacturer’s instructions. DNA extracts were amplified targeting the ITS2 region using the primers ITS-S2F^61^ and ITS-S4R^62^. PCR was carried out in a 25 μL total volume using 12.5 μL of Q5 High-Fidelity 2X Master Mix (New England Biolabs), 1.25 μL of each primer (10 μM), and 1 μL of DNA (10 ng/μL). Reactions were performed in a T100 Thermal Cycler (BioRad^™^) using the temperature profile consisting of an initial denaturation of 98°C for 3 min, followed by 35 cycles of 98°C for 10 s, 52°C for 30 s, and 72°C for 40 s, and a final extension of 72°C for 2 min. The amplicons were Sanger sequenced at STABVIDA Inc. (Portugal) and then analysed using Mega v10.1.7^63^.

**Table 2.** Number of sequences/corresponding taxa that were removed/retained/added by the curation pipeline (automated curation, manual list curation, and manual sequence addition) from/in/to the ITS2 global and crop databases.

| Filters/steps | Global database |  | Crop database |  |
| --- | --- | --- | --- | --- |
|  | Sequences removed | Sequences/taxa retained | Sequences removed | Sequences/taxa retained |
| Start | - | 354,690/119,830 | - | 4,206/81 |
| **Automated curation** |  |  |  |  |
| Fungal sequences | 127 | 354,563 | 3 | 4,203 |
| Non-ITS2 sequences | 29,341 | 325,222 | 249 | 3,954 |
| Incomplete taxonomy | 6 | 325,216 | 0 | 3,954 |
| *Chlorophyta* sequences | 781 | 324,435 | 0 | 3,954 |
| High intraspecific variability | 16,711 | 307,724 | 611 | 3,343 |
| **Total** | **46,966/8,453** | **307,724/111,377** | **863/0** | **3,343/81** |
| **Manual list curation** |  |  |  |  |
| Misidentified sequences | 6 | 307,718 | 0 | 3,343 |
| Taxonomy corrected | 5* | 307,718 | 0 | 3,343 |
| **Total** | **6/3** | **307,718/111,374** | **0/0** | **3,343/81** |
| **Manual sequence addition** |  |  |  |  |
| New sequences | - | 259/182 |  |  |
| **Gran-total** | **-** | **307,977/111,382** | **0/0** | **3,343/81** |

\*Number of detected sequences with incorrect taxonomic classification; These were not removed but instead corrected and retained in the database.

From the 536 samples submitted to DNA sequencing, 259 clean and sufficiently long high-quality sequences were generated, representing 182 species (Table 2). The new sequences were collected in a *fasta* format file and then added to the global database using the manual sequence addition script, as described above. These sequences are also available in the GitHub repository.

#### Country-level databases

After curation, the global ITS2 database was subdivided into two local ITS2 databases for each of the 27 EU countries, according to the local flora retrieved from two online flora databases: Euro+Med PlantBase (https://www.emplantbase.org/home.html) and GBIF (https://www.gbif.org/). These databases complement each other, enabling a more comprehensive representation of the local flora across the 27 EU countries. A more extensive list of plant taxa was retrieved from GBIF than from the Euro+Med PlantBase for the 27 EU countries. Still, there were taxa in the Euro+Med PlantBase list that were missing in the GBIF list.

## Data Records

All final ITS2 databases are publicly available on Zenodo: (i) global database: https://doi.org/10.5281/zenodo.7968519; (ii) crop database: https://doi.org/10.5281/zenodo.7969940, and (iii) country-level databases for the 27 EU countries: https://doi.org/10.5281/zenodo.7970046. The curation scripts are also available at https://github.com/chiras/database-curation.

### Global database

The global database downloaded from GenBank originally held a total of 354,690 sequences, representing 119,830 unique species (Table 2). However, many sequences had problems and were thus removed after the automated implementation of the five sequential curation filters, as follows: (i) 127 fungal sequences; (ii) 29,341 non-ITS2 sequences; (iii) six sequences with incomplete taxonomies; (iv) 781 *Chlorophyta* sequences; and (v) 16,711 sequences with unexpectedly high intraspecific variability for the respective species. After this automated curation, 307,724 sequences (13% loss) were retained in the global database representing 111,377 species (7% loss). The manual list curation detected 11 misidentified sequences in the global database, of which six were removed due to incorrect taxonomic classification, which was not possible to edit, and five were replaced by their correct taxonomic classification. After this additional step, a total of 307,718 sequences, representing 111,374 species, were retained in the global database. With the addition of our own ITS2 sequences, the final global database contains 307,977 sequences, representing 534 families, 11,034 genera, and 111,382 species of vascular plants.

### Crop database

A list of European crop species, containing for each entry an accurate taxonomic classification string, was carefully assembled and then used to retrieve the matching sequences from GenBank. A total of 4,206 sequences, representing 81 taxa, were downloaded from GenBank. The automated curation workflow identified and removed from this database the following number of sequences: (i) three fungal sequences; (ii) 249 non-ITS2 sequences; and (iii) 611 sequences with high intraspecific variability for the respective species (Table 2). As expected from the nature of the assembled list, no ‘Incomplete taxonomy’ or ‘*Chlorophyta*’ problems were detected. Furthermore, no sequences were removed or added by the ‘Manual list curation’ and ‘Manual sequence addition’ components of the pipeline. Accordingly, the final crop database comprises 3,343 sequences (21% loss), representing 25 families, 50 genera, and 81 species (0% loss).

### Country-level databases

Table 3 compiles the sizes of the two ITS2 databases generated for each of the 27 EU countries, by taking into account the local floras extracted from Euro+Med PlantBase and GBIF. The 27 ITS2 databases generated using the Euro+Med PlantBase lists cover between 66% and 89% of the vascular plant species listed for each country (Fig. 4). The ITS2 databases of the Mediterranean countries show the lowest coverage of the local flora, with Greece having 66%, Spain 69%, France 71%, and Italy 72%. In contrast, the ITS2 databases obtained for the Baltic countries contain sequences representing a high proportion of their plant diversity, with Latvia (89%) at the top of the ranking, followed by Lithuania and Estonia (88%), and Finland (87%). The findings for Mediterranean countries were expected due to their higher species richness, thereby requiring a higher sequencing effort to achieve the levels of the Baltic countries. Apart from Malta, the lists extracted from GBIF are species-richer than those extracted from Euro+Med PlantBase, explaining the lower coverage of the corresponding ITS2 databases. Hence, the coverage of the ITS2 databases generated using the GBIF lists is lower than that generated using the Euro+Med PlantBase lists, varying between 31% for France and 86% for Lithuania (Fig. 4).

**Table 3.** Sizes of the country-level ITS2 databases in relation to the vascular plant species inventories extracted from (A) Euro+Med PlantBase (https://www.emplantbase.org/home.html) and (B) GBIF platforms (https://www.gbif.org/). Number of ITS2 sequences, number of species with ITS2 sequences, number of species extracted from Euro+Med PlantBase and GBIF, and proportion of species with ITS2 sequences in the database.

| Country | Sequences<br>A/B | Species with<br>sequences<br>A/B | Species in the<br>flora<br>A/B | Species<br>coverage (%)<br>A/B |
| --- | --- | --- | --- | --- |
| Austria | 25,209/40,297 | 2,747/5,141 | 3,572/11,316 | 77/45 |
| Belgium | 18,083/57,279 | 1,810/9,070 | 2,182/17,359 | 83/52 |
| Bulgaria | 23,460/26,591 | 2,812/3,137 | 3,839/5,599 | 73/56 |
| Croatia | 20,640/28,011 | 2,400/3,385 | 3,053/5,852 | 79/58 |
| Cyprus | 10,777/13,524 | 1,296/1,624 | 1,710/2,600 | 76/62 |
| Czechia | 18,620/32,640 | 1,929/3,707 | 2,411/6,650 | 80/56 |
| Denmark | 16,804/29,735 | 1,583/3,218 | 1,846/5,307 | 86/61 |
| Estonia | 14,709/29,816 | 1,352/3,363 | 1,534/5,586 | 88/60 |
| Finland | 16,214/27,716 | 1,527/2,958 | 1,747/5,351 | 87/55 |
| France | 32,207/59,623 | 4,134/9,426 | 5,799/30,227 | 71/31 |
| Germany | 26,709/57,740 | 2,979/8,516 | 4,107/20,101 | 73/42 |
| Greece | 24,909/34,231 | 3,550/5,033 | 5,382/10,966 | 66/46 |
| Hungary | 20,670/30,557 | 2,089/3,584 | 2,535/6,925 | 82/52 |
| Ireland | 14,341/18,944 | 1,328/1,884 | 1,582/2,682 | 84/70 |
| Italy | 32,955/47,441 | 4,310/6,808 | 5,948/16,427 | 72/41 |
| Latvia | 13,405/21,771 | 1,210/2,178 | 1,367/3,148 | 89/69 |
| Lithuania | 14,504/16,702 | 1,316/1,486 | 1,497/1,736 | 88/86 |
| Luxembourg | 1,627/22,033 | 137/2,147 | 178/3,085 | 77/70 |
| Malta | 9,874/6,672 | 1,108/677 | 1,371/929 | 81/73 |
| The Netherlands | 16,693/43,884 | 1,530/5,770 | 1,881/11,115 | 81/52 |
| Poland | 22,099/33,094 | 2,256/3,711 | 2,785/6,740 | 81/55 |
| Portugal | 19,593/37,654 | 2,372/5,103 | 3,031/10,347 | 78/49 |
| Romania | 25,349/28,296 | 2,820/3,221 | 3,673/5,858 | 77/55 |
| Slovakia | 18,344/26,330 | 1,925/2,795 | 2,448/4,616 | 79/61 |
| Slovenia | 18,329/25,063 | 1,968/2,727 | 2,434/4,129 | 81/66 |
| Spain | 32,265/58,332 | 4,384/9,628 | 6,380/27,128 | 69/35 |
| Sweden | 20,468/48,305 | 2,048/6,736 | 2,446/14,648 | 84/46 |

**Fig. 4.**
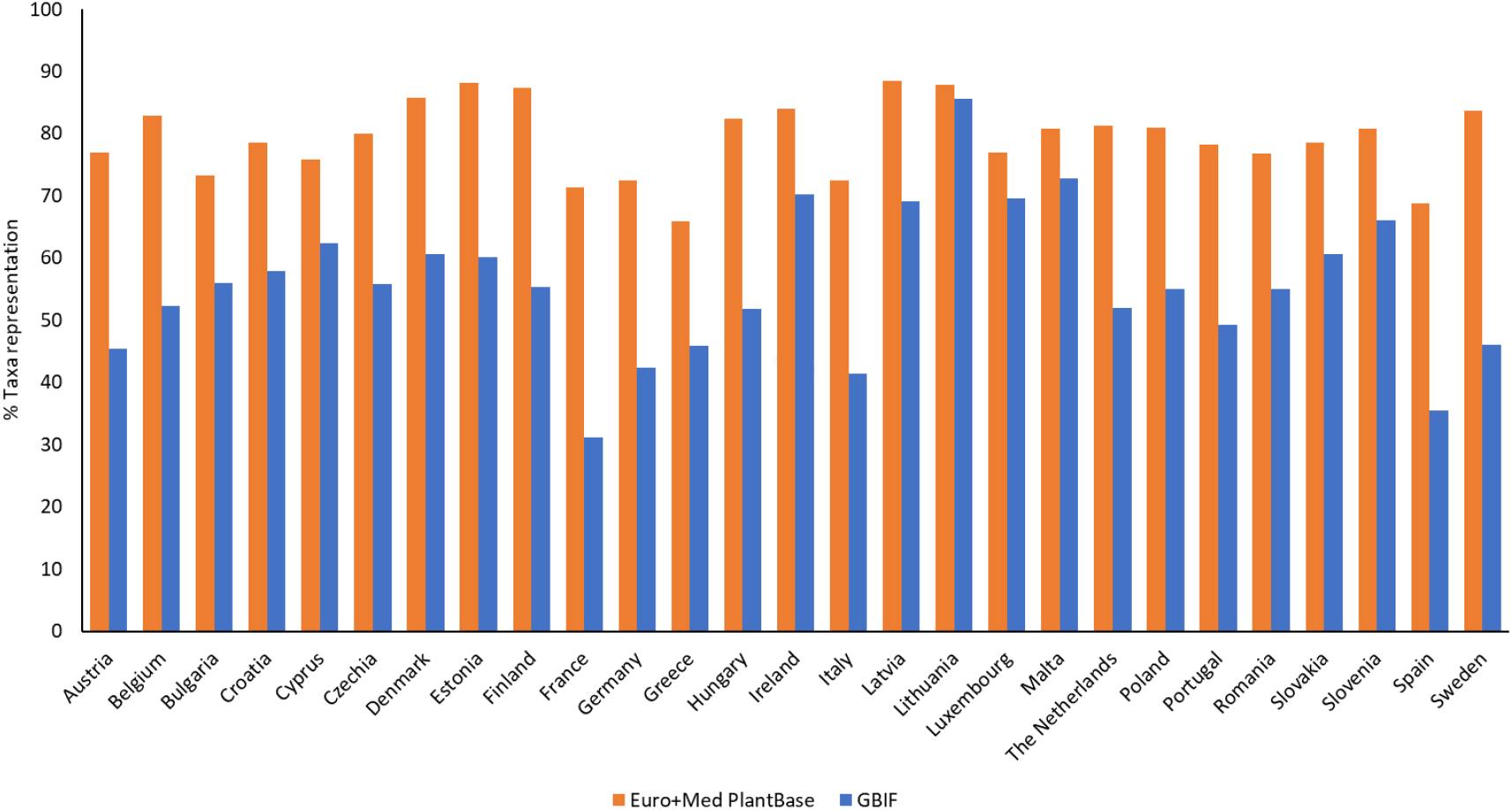
Taxa representation of the two reference ITS2 databases generated for each of the 27 EU countries, using the flora information extracted from the Euro+Med PlantBase (https://www.emplantbase.org/home.html) and GBIF platforms (https://www.gbif.org/).

**Fig. 4.**
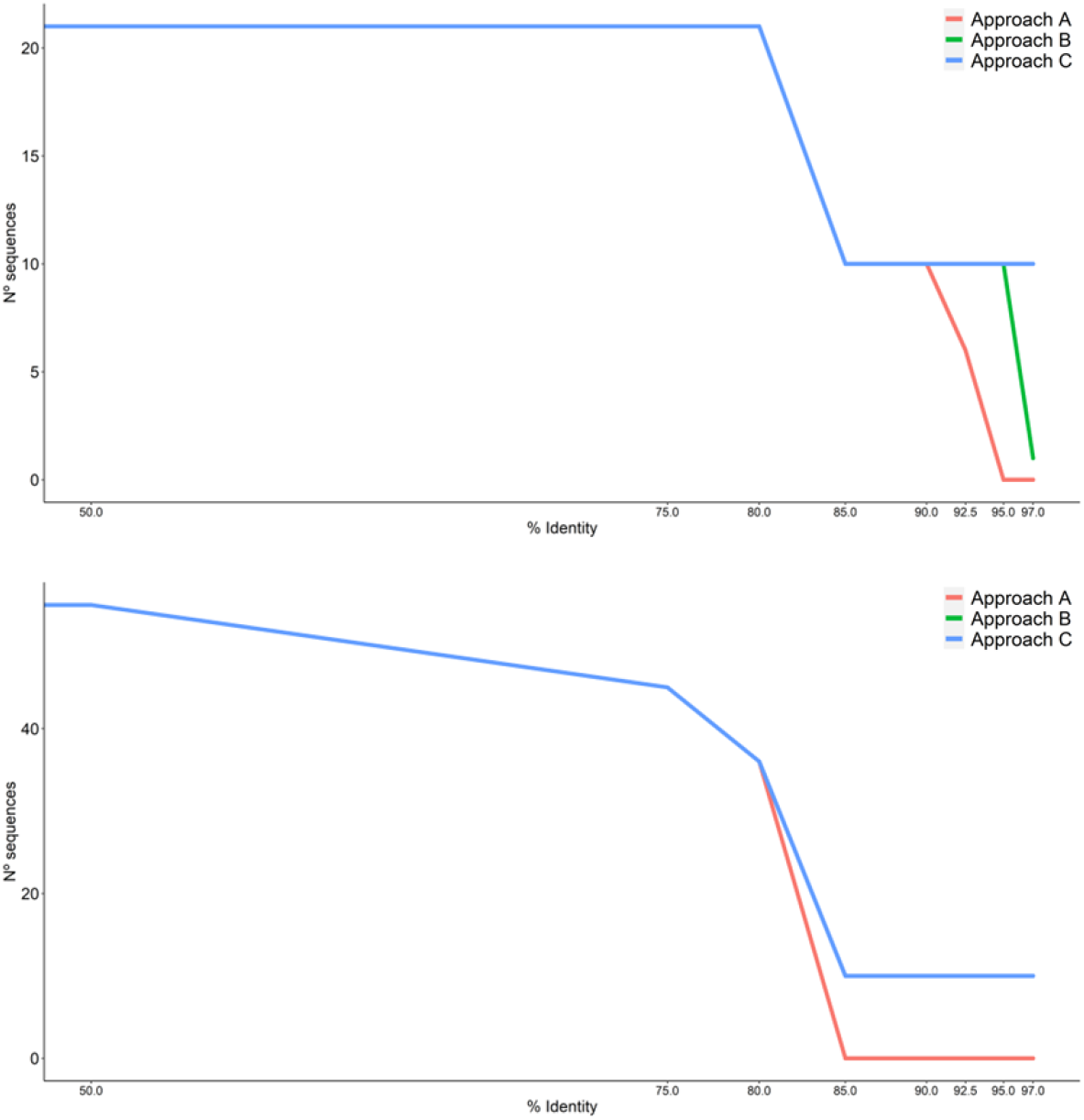
Number of sequences retained in the ITS2 database for *Malus pumila* (top chart) *and Pyrus communis* (bottom chart) by the automated curation workflow. Approach A: sequences with a median identity < 97% in pairwise all-against-all global alignments are removed in a single iteration; Approach B: sequences are removed iteratively using an incremental drop-out identity threshold of 50%, 75%, 80%, 85%, 90%, 92.5%, 95%, and 97%; Approach C: sequences are removed using the incremental threshold of ‘Approach B’ while ensuring that 50% of the initial sequences are retained in the database.

## Technical Validation

### Fungal sequences identified as plants in GenBank

A total of 127 fungal sequences were detected among the sequences identified as plants in GenBank. Of these, 55 (43%) belonged to the phylum *Ascomycota*, and the most common genera were *Erysiphe* (15%), *Aspergillus* (14%), *Davidiella* (11%), *Gibberella*, and *Mycosphaerella (8%)*, and *Eurotium* (6%). These fungi are either pathogens or endophytes commonly detected in plant tissues (e.g., *Erysiphe* causes powdery mildew, and *Mycosphaerella* causes leaf blight). Fungal PCR-amplifications from infected plant tissues are well documented for ITS2 primers designed for plants^64^, explaining the misidentified sequences deposited in GenBank. One such example comes from the single ITS2 sequence available in GenBank for *Rumex stenophyllus* (accession number MG235257). During the automated curation, this sequence was identified as belonging to the genus *Alternaria*, leading to its removal from the global database.

### Plant sequences assigned an incorrect taxonomic classification

The automated curation allowed identification and removal of sequences that were deposited in GenBank with incorrect taxonomic classification. For instance, the sequences with accession numbers KF454376 and KF454377, originally identified in GenBank as *Typha angustifolia* (*Typhaceae*), turned out to belong to the genus *Taraxacum* (*Asteraceae*) after manual verification. With the intraspecific analysis implemented by the fifth filter of the automated curation, these two sequences were automatically removed from the global database.

### Assessment of intraspecific variability

The accuracy of the taxonomic classification depends on the power of the chosen marker in discriminating between interspecific and intraspecific variation, i.e., the overlap of the genetic variation between species should be small or ideally non-existent. Hybridization is a common natural or human-mediated phenomenon in many wild plant species as well as in many crops, such as *Brassica napus* and *Brassica rapa*, or *Malus domestica* and *Pyrus communis*. This erodes species delimitations and increases intraspecific variability, making automated curation a more challenging endeavour.

The last step of the automated curation (the fifth filter) applies a deterministic assessment of intraspecific variability for the respective species. In the initial configuration of the pipeline, the sequences that had a median identity lower than 97% in pairwise all-against-all global alignments were removed from the database in a single iteration. This revealed itself to be very stringent for taxa suffering from high intraspecific variability, leading to the removal of all the sequences from the curated database. Hence, this direct approach (approach A) was replaced by the iterative increment of the drop-out threshold (approach B), as explained in the section ‘Automated curation’. While an improvement in the pipeline’s performance was noted, there was still a low number of retained sequences in the curated database (e.g., *Malus domestica* was represented by a single sequence). Lastly, in the final configuration of the automated curation (see the ‘Automated curation’ section), the introduced threshold that retains 50% of the initial sequences (approach C) seems to represent a good trade-off between removing taxa with wrong GenBank taxonomic assignments and retaining the sequences that are still within expected intraspecific variability.

The outcomes of these three approaches are illustrated in Fig. 5 for *Malus pumila* and *Pyrus communis*. No sequences or a single sequence were retained in the curated database for *Malus pumila* with approaches A and B, respectively. In contrast, 10 of the initial 20 sequences were retained in the curated database at 85% identity when approach C was applied. In the case of *Pyrus communis*, approaches B and C performed equally well, retaining 10of the initial 55 sequences, whereas all the sequences were removed from the database when applying approach A.

**Fig. 5.** Number of publications for each of the *Viridiplantae* DNA barcoding markers, used singly or in combination in DNA metabarcoding studies. Information retrieved from Web of Science, accessed on 22 February 2023, using the following search strings: “ITS2 (All Fields) AND metabarcoding (All Fields) NOT fung* (All Fields) NOT diatoms (All Fields)”; “rbcL (All Fields) AND metabarcoding (All Fields) NOT fung* (All Fields) NOT diatoms (All Fields)”; “trnH-psbA OR psbA-trnH (All Fields) AND metabarcoding (All Fields) NOT fung* (All Fields) NOT diatoms (All Fields)”; “matK (All Fields) AND metabarcoding (All Fields) NOT fung* (All Fields) NOT diatoms (All Fields)”. The retrievals were manually verified and the publications that were not targeted to plants were ignored.

**Fig. 6.** Number of publications on ITS2 metabarcoding in the last ten years. Information retrieved from Web of Science, accessed on 22 February 2023, using the search string “ITS2 (All Fields) AND metabarcoding (All Fields) NOT fung* (All Fields) NOT diatoms (All Fields)”.

**Fig. 7.** Schematic representation of the curation pipeline. The components ‘Automated curation’, ‘Manual list curation’, and ‘Manual sequence addition’ can be used singly or in conjunction.

**Fig. 8.** Taxa representation of the two reference ITS2 databases generated for each of the 27 EU countries, using the flora information extracted from the Euro+Med PlantBase (https://www.emplantbase.org/home.html) and GBIF platforms (https://www.gbif.org/).

**Fig. 9.** Number of sequences retained in the ITS2 database for *Malus pumila* (top chart) *and Pyrus communis* (bottom chart) by the automated curation workflow. Approach A: sequences with a median identity < 97% in pairwise all-against-all global alignments are removed in a single iteration; Approach B: sequences are removed iteratively using an incremental drop-out identity threshold of 50%, 75%, 80%, 85%, 90%, 92.5%, 95%, and 97%; Approach C: sequences are removed using the incremental threshold of ‘Approach B’ while ensuring that 50% of the initial sequences are retained in the database.

### Comparison with other databases

The global ITS2 database generated in this study contains sequences from 111,377 species, representing an increase of over 62% when compared to the databases of Sickel et al.^51^ (72,325 species) and Dubois et al.^12^ (∼70,000 species). The implementation of the automated curation script developed herein is able to resolve troublesome sequences downloaded from GenBank while still retaining a good representation of worldwide species in the curated database. Moreover, the manual list curation step prevents reliable sequences from being removed at the same time that the manual sequence addition step facilitates database enrichment.

## Code Availability

All code used in this study is freely available in https://github.com/chiras/database-curation. The developed global and country-level databases are also provided in the same repository as well as in Zenodo.

## Acknowledgements

AQ acknowledges the PhD scholarship (2020.05155.BD), funded by the Portuguese Foundation for Science and Technology (FCT). This work was developed in the framework of INSIGNIA – Environmental monitoring of pesticide use through honeybees (SANTE/E4/SI2.788418-SI2.788452-INSIGINIA-PP-1-1-2018) and INSIGNIA-EU − Preparatory action for monitoring of environmental pollution using honey bees (Procurement procedure ENV/2021/OP/0014 of 28-09-2021). FCT provided financial support by national funds (FCT/MCTES) to CIMO (UIDB/00690/2020 and UIDP/00690/2020) and SusTEC (LA/P/0007/2021).

## Author contributions

AQ, AK, and MAP conceived the ideas and designed the methodology. AK and AQ developed the scripts and the databases. JR extracted the list of European flora from Euro+Med PlantBase and assisted with the computational resources. Plant leaves for the manual sequence addition were provided by MAP, JA, RB, VB, KG, FH, OK, MP, IR, and FV. AQ, CAYG and MH performed the DNA extractions of the plant leaves. AQ, MAP, and AK, wrote the manuscript. JvdS acquired INSIGNIA’s funding. All the authors critically reviewed the manuscript for important intellectual content.

## Competing interests

We declare no competing interests.

## Notes

https://github.com/chiras/database-curation/tree/main/curation

https://doi.org/10.5281/zenodo.7969940

https://doi.org/10.5281/zenodo.7970046

https://doi.org/10.5281/zenodo.7968519

### Competing Interest Statement

The authors have declared no competing interest.

